# MSAC: Compression of multiple sequence alignment files

**DOI:** 10.1101/240341

**Authors:** Sebastian Deorowicz, Joanna Walczyszyn, Agnieszka Debudaj-Grabysz

## Abstract

**Motivation:** Bioinformatics databases grow rapidly and achieve values hardly to imagine a decade ago. Among numerous bioinformatics processes generating hundreds of GB is multiple sequence alignments of protein families. Its largest database, i.e., Pfam, consumes 40–230 GB, depending of the variant. Storage and transfer of such massive data has become a challenge.

**Results:** We propose a novel compression algorithm, MSAC (Multiple Sequence Alignment Compressor), designed especially for aligned data. It is based on a generalisation of the positional Burrows–Wheeler transform for non-binary alphabets. MSAC handles FASTA, as well as Stockholm files. It offers up to six times better compression ratio than other commonly used compressors, i.e., gzip. Performed experiments resulted in an analysis of the influence of a protein family size on the compression ratio.

**Availability:** MSAC is available for free at https://github.com/refresh-bio/msac and http://sun.aei.polsl.pl/REFRESH/msac.

**Supplementary material:** Supplementary data are available at the publisher Web site.

## 1 Introduction

In modern times, the sizes of data collected in various fields are huge and are constantly growing. This has a direct impact on the increasing costs of IT infrastructure. As far as bioinformatics is concerned a number of surveys claim that soon the cost related to storage of data will be at least comparable to the costs of generating these data (Deorowicz & Grabowski, 2013; Stephens *et al*., 2015). A natural solution to this problem is to reconsider the need to store all collected data. One could advocate that a significant part of collected files are rather “temporary” and can be safely removed after performing the analyses. Unfortunately, this could ruin one of the fundamentals of science, i.e., reproducibility, as many bioinformatics tools work in a non-deterministic manner. That means that when using these tools several times, we would obtain similar results, but not always exactly the same. What is more, the emergence of new bioinformatics software allows revealing undiscovered relationships between collected data and their new biological meaning. Hence, very often the reproducibility is of high importance, so a number of temporary files are stored for a long time in many research centres.

To make storage possible compressing data seems to be a primary solution. Even 25-years old, but still popular gzip algorithm, treated as de facto standard, allows to reduce many types of data several times. That is why gzip is a common choice in bioinformatics when storing sequencing data, protein sequences, etc. Nevertheless, gzip is a universal compressor, developed mainly to compress textual files with sizes considered to be small from the modern perspective. Besides, being all-purpose, it cannot make use of special types of redundancy existing in the bioinformatic datasets. Therefore, a number of specialised compressors were developed in recent years.

The most attention was devoted to the field of genome sequencing (Bonfield & Mahoney, 2013; Roguski & Deorowicz, 2014; Numanagić *et al*., 2016). The obtained compression ratios are sometimes an order of magnitude better than offered by gzip. Other examples of huge datasets whose storage is supported by dedicated software are collections of complete genomes of the same species. Various algorithms for reducing their sizes a few orders of magnitude were proposed (Deorowicz *et al*., 2013; Li, 2016). In addition, these algorithms may have offered fast queries to the compressed data. Results of ChiP-Seq and RNA-seq experiments are another noteworthy example. Solutions capable of compressing them by the factor of at least a few have been proposed (Wang *et al*., 2016; Ravanmehr *et al*., 2017) very recently.

Storage of protein databases, like Pfam (Fin *et al*., 2016), is also a challenge. For example, the recent release of Pfam (v. 31.0) contains 16,712 protein families and about 5 billion residues. The uncompressed size of the default variant of the database is about 42 GB. Moreover, its NCBI variant stores more than 22 billion residues and occupies about 228 GB. Even when gzip-compressed, the files are still of extraordinary sizes, i.e., 5.6GB and 24GB, respectively.

To the best of our knowledge, there has been no breakthrough in the field of compression of such protein datasets so far. An interesting alternative could be to use a better universal compressor and the popular 7-zip program seems to be a reasonable choice. Nevertheless, it consumes a lot of memory for large files and is relatively slow. What is more important, it is still a universal tool, thus it does not use knowledge about properties of these datasets. Therefore, significantly better results should be possible to obtain if we took this information into account and use it during the construction of the algorithm.

In this article, we propose a specialised compression algorithm designed to decrease a size of databases of multiple sequence alignments (MSA) of protein families. Our tool, MSAC, can compress both FASTA files containing a single protein family data as well as collections of proteins in Stockholm format (Fin *et al*., 2016) (used in Pfam database). The proposed algorithm not only offers several times better compression ratios than gzip (and significantly better than 7-zip) for large protein families. It is also much faster in compression than the examined counterparts and has moderate memory requirements, roughly similar to the input dataset (or a single family for Stockholm files) size.

## 2 Methods

### 2.1 Background

The heart of gzip and 7-zip compressors is a well known Ziv–Lempel family of algorithms (Ziv & Lempel, 1977; Storer & Szymanski, 1982). Its key idea is to look for exact copies of some parts of a file being compressed. When a copy is sufficiently long (usually at least 3 or 4 symbols), then it is beneficial to store the information about its location and its length instead of storing its symbols literary. Roughly speaking, both gzip and 7-zip perform so-called LZ-factoring in which the input sequence of symbols is transformed into a sequence of tuples that can be of two types: matches (describing a repetition of a fragment of already processed part of the file) and literals (single symbols stored when no sufficiently-long match was found). This sequence of tuples is then coded using one of the entropy coders like Huffman coder (Huffman, 1952) or range coder (Salomon & Motta, 2010). In case of MSA files, looking for copies of the fragments of a text means looking for identical fragments of sequences of various proteins. In this way long runs of gaps, that supplement alignments, are stored as copies of some other “lines” of MSA files. Such a strategy leads to up to 15-fold compression in case of gzip and up to 50-fold for 7-zip, when compressing large protein families with many gaps.

When considering MSA files several drawbacks of the mentioned approach can be shown. The first one results from the specificity of an index structure, which is necessary to search for matches in the part of the file that has been just processed. Such index structures (like suffix trees, hash tables) are usually several times larger than the indexed part of the data. The second disadvantage results from the dependence between the size of the indexed part and the number of positions that should be examined when looking for the best match. Asa consequence, the operating time increases together with the growth of the indexed part. To preserve speed, LZ-compressors have to make a trade-off, e.g., restrict to examining only some fraction of potential matches. What’s more, defining what “the best match” means is not clear. Coding of the longest possible match not always leads to the highest compression ratios. It happens that choosing a shorter match (or even a literal) at a current position leads to a longer match at the following position, which in consequence improves the compression ratio.

Besides, in case of MSA files, the sequences are aligned along the columns and it is unusual to find a long match starting at different column than the current one. That is why, most of potential candidates for matches could be omitted during analyses. Finally, this knowledge (inaccessible to universal compressors) could be used to encode the found matches cheaper, i. e, using the smaller number of bits.

In the universal compression field there are also different approaches than LZ-based. The two most known families are based on the Burrows–Wheeler transform (BWT) and prediction by partial matching (PPM) (Cleary & Witten, 1984). The popular bzip2 program, implementing Fenwick’s variant (Fenwick, 1996) of the BWT-based compressor, is a representative of the former family. The Burrows–Wheeler transformation (Burrows & Wheeler, 1994) of the input sequence is performed at its first stage. As a result, long parts of the transformed sequence contain only a few different symbols (or even a single symbol). Such a temporary sequence is then further changed using a move-to-front transformation (Bentley *et al*., 1986) and Huffman encoded (Huffman, 1952). In the PPM algorithms the statistics of symbol occurrences in various contexts (formed by preceding symbols) are collected. They are used to estimate the probability of appearance of each symbol in every single place. Based on this estimation short codewords are assigned to more probable symbols and longer ones to less probable, with the use of an entropy coder. An interested reader is referred to one of textbooks on data compression, for more detailed discussion and examples of the LZ-based, BWT-based, and PPM-based algorithms (Salomon & Motta, 2010).

### 2.2 General idea of the proposed algorithm

To overcome the problems with indexing of huge files and make use of the alignment property of the MSA files we decided not to follow the obvious strategy of implementing the LZ-based algorithms adopted for MSA data. Rather than that, we focused on the recently proposed, positional Burrows–Wheeler transform (PBWT) (Durbin, 2014). Its name reflects that it was motivated by the classical Burrows–Wheller transform. Nevertheless, it was designed to transform aligned bit vectors to allow faster queries for genotype data. Later, Li used the PBWT to develop a specialised compressor of genotype datasets (Li, 2016). One of the assets of the PBWT is its memory frugality as no large index structures (required not only by LZ-based algorithms, but also by classical BWT-based and PPM-based ones) are necessary. Instead, the original PBWT processes rather short bit vectors (of values 0 and 1) one by one.

The PBWT was defined for bit vectors, but fortunately it can be simply generalized to larger alphabets. In the next section we propose such a generalisation. Then, we adopt some transforms known from the BWT-based compressors to obtain the novel algorithm for the MSA data.

### 2.3 Positional Burrows–Wheeler transform for non-binary alphabets

Let Σ be an alphabet of symbols {0, 1,…, *σ −* 1}. Let *S* be an ordered collection of equal-length sequences {*S*^1^, *S*^2^,…, *S^n^*}. The length of each sequence is denoted by *ℓ* and for each valid *i*: 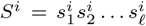. Each 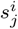 is a symbol from the alphabet Σ. A substring of *S^i^* from *j*th to *k*th symbols is defined as 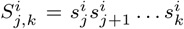. A special type of a substring is a suffix of a sequence: 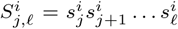. The *S* can be seen as a matrix of *n* rows and *ℓ* columns. For simplicity of presentation it is convenient to define also a *j*th column of *S* as 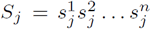. Finally, by sort*_j_*, (*S*) we denote the collection *S* lexicographically sorted according to the suffixes starting at (*j* + 1)th symbol.

The *generalised positional Burrows–Wheeler transform* (gPBWT) changes *S* into *P*, where *P* is defined in the following way. The last column of *P* is equal to the last column of *S*, i.e., *P_ℓ_* = *S*_ℓ_. To obtain *P_j_* we sort the sequences of *S* according to their suffixes starting from the (*j* + 1)th symbol and pick the *j*th column, i.e., *P_j_* = (sort*_j_* (*S*))*_j_*.

The original PBWT by Durbin transforms bit vectors with the prefix-sorting order. We decided to present the gPBWT algorithm in the suffix-sorting order just to be more similar to the original BWT definition. It is worth to mention that both orderings for PBWT and gPBWT are equivalent in the sense that it is enough to reverse the input sequences to switch between the variants.

What is important from the performance point of view, the sort*_j_* (*S*) can be easily obtained from sort*_j_*_+1_(*S*). Moreover, in practice it suffices to store the ordering of indices of the *S* sequences and no coping of the complete sequences of *S* is made. The pseudocodes of algorithms for determination of the gPBWT and its reverse are presented in Fig. 1. In the gPWBT construction, for the jth column, we calculate the histogram *H* of symbols at the *j*th column. Then we compute the cumulative statistics *H\**, where *H\**[*c*] stores the information about the total number of symbols lexicographically smaller than *c* in the *j*th column. In the next step, we make use of the ordering of suffixes obtained when processing (*j* + 1)th column and *H\** to obtain the ordering according to the suffixes starting from the *j*th column. Finally, we permute the symbols of the *j*th column of *S* to obtain the *j*th symbol of *P* using the ordering according to suffixes of *S* starting form the (*j* + 1)th symbol. The reverse gPBWT algorithm performs essentially the same steps, but some of them in the opposite order.

**Fig. 1.**
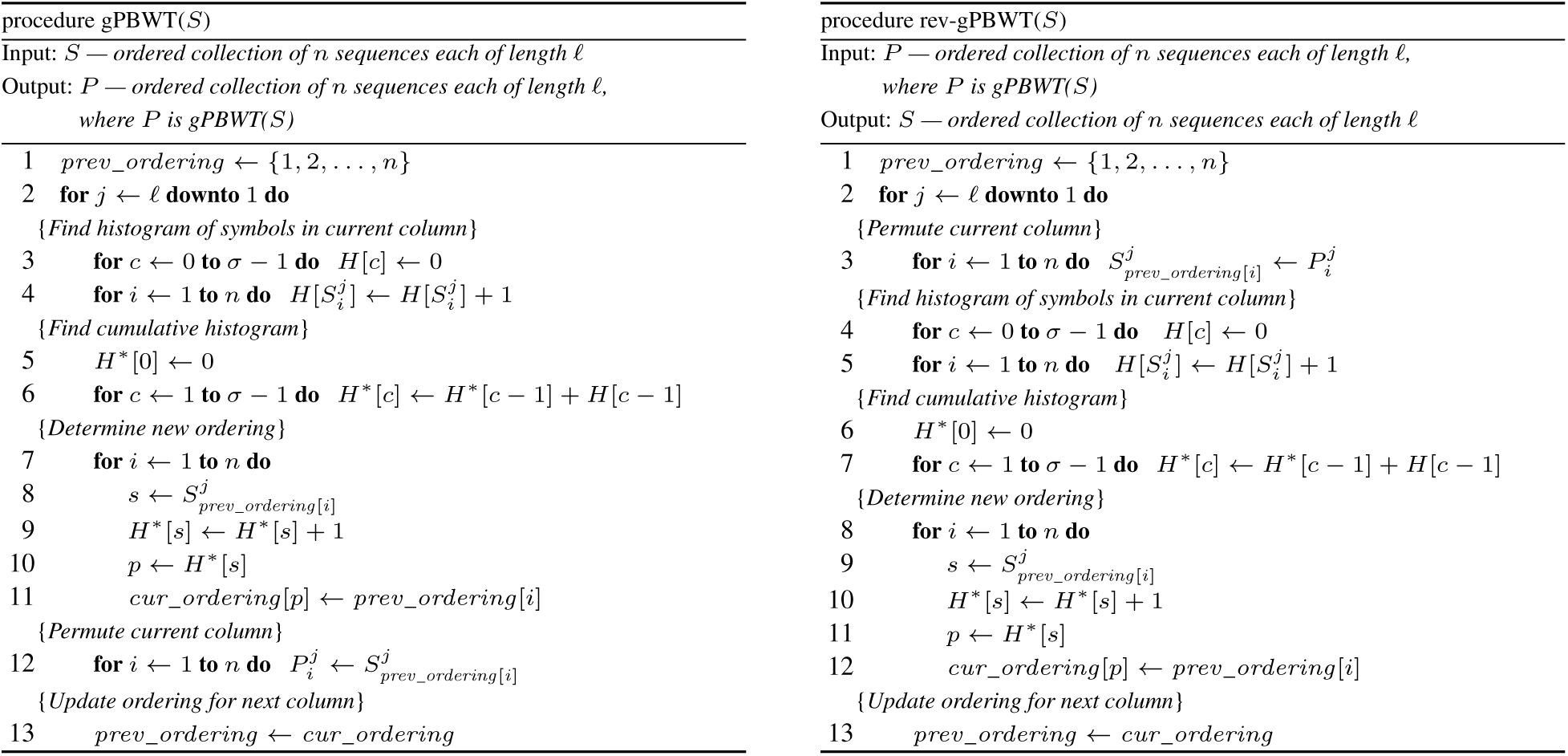
Pseudocodes of the generalised PBWT algorithm (left) and its reverse (right)

Let us focus now on the time and space complexity of the gPWBT computation. For each column we have to initialize array *H* and calculate array *H\** which is made in *O*(σ) time. Determination of the new ordering, as well as performing the permutation takes *O*(*n*) time. Since there are *ℓ* columns, the total time complexity of both the forward as well as the reverse gPBWT algorithms is *O*(max(*n*, *σ*), *ℓ*). In case of sufficiently large collections *S*, i.e., when the number of sequences *n* is not smaller than the alphabet size *σ*, the time complexity reduces to *O*(*nℓ*), which is linear in terms of the total number of symbols in the input dataset *S*. Since the only data that must be stored in memory consists of the previous and current orderings and the statistics *H* and *H\**, the space complexity of the algorithm is just *O*(max(*n, σ*)), which for sufficiently large collections is just *O*(*n*).

### 2.4 Novel compression algorithm

The compression algorithm we propose in the article is able to handle both FASTA and Stockholm files. FASTA files contain a single (aligned) protein family whereas Stockholm files can be seen as a concatenation of alignments of many protein families supplemented by some metadata. For simplicity we start with the description of the variant for FASTA files.

The general scheme of the proposed compression algorithm is presented in Fig. 2. In the first stage, the input FASTA file is read into the memory (Read-and-Split block). The ids of the sequences are concatenated into a single string to be transferred to the LZMA block where they are compressed using the LZMA algorithm (used for example in the 7-zip program). The raw protein sequences are transferred to the Transpose block for the transposition of the matrix (containing the protein sequences in the form of rows). The columns of the matrix S (rows of the transposed S) are transferred to the gPBWT block from the last to the first one. In this block they are transformed by the gPBWT algorithm described in the previous subsection.

**Fig. 2.**
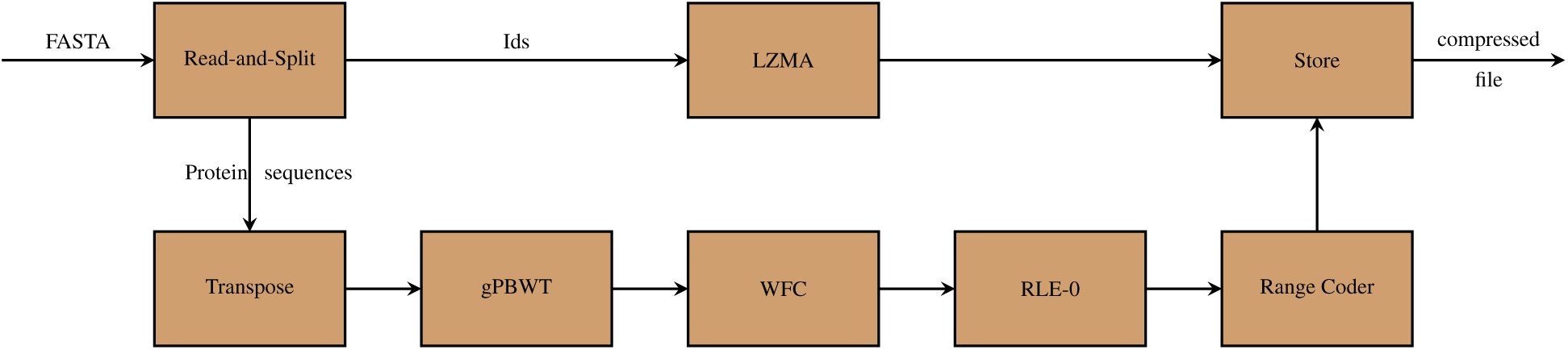
Scheme of the proposed compression algorithm

Each permuted column is then transferred separately to the WFC block. This block implements the *weighted frequency count* transform (WFC) (Deorowicz, 2002). The WFC transform changes the column of symbols over any alphabet into a column of integers. Roughly speaking, for each position of the column which is transformed, WFC predicts a rank for each alphabet symbol (the smaller the rank, the more frequently the alphabet symbol appeared in the former part of the column). Then the current symbol is replaced by its rank. The output of this block is a column of integers. The integers are usually small and majority of them (sometimes even up to 90 percent) are zeroes.

In the next stage the zero-run-length-encoding transform (Fenwick, 1996a), RLE-0 block, replaces the repetitions of zeroes using a simple coding scheme: 0 is replaced by 0*_a_*, 00 → 0*b*, 000 → 0*_a_*0*_a_*, 0000 → 0*_a_*0*_b_*, 00000 → 0*_b_*0*_a_*, 000000 → 0*_b_*0*_b_*, 0000000 → 0*_a_*0*_a_*0*_a_*, etc. This reduces the length of columns noticeably.

In the next stage, the column is entropy coded. For this purpose we use a *range coder* (Salomon & Motta, 2010) (RangeCoder block). This coder assigns short codewords to frequent symbols and longer to the rare ones, which results in significant reduction of the space necessary to represent the input sequence. Since the frequency of symbols in the input of this block could differ by a few orders of magnitude (which causes some problems to entropy coders), we employ a simple modelling. For each symbol we initially encode a flag indicating whether it is 0*_a_*, 0*_b_*, 1, or something larger. In most cases we are ready here, but when the symbol belongs to the last group (it is larger than 1), we proceed in the following way. We encode the group id of the symbol, where the available groups are: {2, 3}, {4,…, 7}, {8,…, 15}, {16,…, 31}, {32,…, 63}. Then we encode the value of the symbol withing a group. The maximal value of the integer (63) reflects the fact that the allowed symbols in the input sequences are lower- and upper-case letters, and a few special values, i.e., ‘−’, ‘.’, ‘*’.

To improve the compression ratio even more, the symbols within a group are encoded in contexts, i.e., the probability of appearance of each symbol (necessary for range coder) is estimated taking into account a short-time history. For example, the context for the flags is defined by up to 5 recently encoded flags. The contexts for group ids and symbols inside groups are composed from up to 3 recently encoded group ids. Finally the compressed sequences for each column and the LZMA-compressed ids are collected in a single output file (Store block).

The compression of Stockholm files containing alignments of many protein families is similar. The file is processed in parts where each part contains a single family. At the beginning the family data are split into metadata and alignment data. The metadata are LZMA compressed while the alignment data are compressed using the algorithm for the FASTA files.

Our algorithm was implemented in the C++14 programming language. To make use of the multi-core architecture of modern CPUs, it is implemented using the C++ native threads. The main thread is responsible for Read-and-Split and Store stages, as well as it controls the execution. The stages Transpose, gPBWT, RLE-0, and RangeCoder are made by their own threads. As WFC is the most time consuming, it is executed by up to 4 separate threads (each of them independently processes some subset of columns). The software was compiled using GCC 6.2 with -O3 optimization enabled.

## 3 Results

The test platform was equipped with two Intel Xeon E5–2670 v3 CPUs (clocked at 2.3 GHz, 24 physical cores in total) and 128 GiB of RAM. In the experiments we limited the number of threads used by the compressors to four. As the test datasets we picked the default Pfam v. 31.0 collection of protein sequences (Fin *et al*., 2016). The Stockholm file with the whole collection has size of 41.6GB and contains 16,479 protein families. To examine the scalability of our software by the growing sizes of collections, we evaluated also the larger variants of Pfam database: Uniprot and NCBI. They are two and five times larger, respectively, than the default collection.

Currently, gzip is the most commonly used compressor for MSA data, so it was an obvious choice as the benchmark. Alas, gzip is a single-threaded application, so for a fair comparison we used its multi-threaded variant, i.e., pigz. As, to the best of our knowledge, there are no specialised compressors for MSA files, we examined one more universal tool, namely 7-zip. It is quite often used when the better compression ratio is required, and the slower compression and decompression speeds could be accepted.

MSAC was designed for both separate FASTA files (each file containing one protein family data) and collections stored in Stockholm files. Initially we evaluated the compression of the whole Stockholm files. The results given in Table 1 show that our compressor achieves the compression ratio (defined as the size of the original file divided by the size of the compressed file, so the higher the value, the better) about 24, which is 3.5 times better than the ratio offered by gzip. In real numbers, the default Pfam collection can be stored in as little as 1.74 GB (compared to 5.6 GB for gzip). MSAC compression ratio is also 37 percent better than that of 7zip. Similar observations can be made for Uniprot and NCBI variants of Pfam database.

**Table 1.** Compression results for complete collections of Pfam database.

| Dataset | No. of families | No. of sequences | File size | pigz |  |  | 7zip |  |  | MSAC |  |  |
| --- | --- | --- | --- | --- | --- | --- | --- | --- | --- | --- | --- | --- |
|  |  |  |  | size | c-time | d-time | size | c-time | d-time | size | c-time | d-time |
| Stockholm files |  |  |  |  |  |  |  |  |  |  |  |  |
| Pfam-A | 16,479 | 31,051,470 | 41.6 | 5.60 | 4,723 | 227 | 2.38 | 6,130 | 379 | 1.74 | 2,766 | 874 |
| Pfam-Uniprot | 16,712 | 84,689,547 | 82.3 | 8.73 | 9,504 | 404 | 3.30 | 13,968 | 632 | 2.63 | 3,600 | 1,904 |
| Pfam-NCBI | 16,712 | 177,952,603 | 227.7 | 26.93 | 24,413 | 1,185 | 10.35 | 41,329 | 1,811 | 8.82 | 28,486 | 4,723 |
| FASTA files |  |  |  |  |  |  |  |  |  |  |  |  |
| Pfam-A | 16,479 | 31,051,470 | 31.9 | 4.83 | 4,495 | 132 | 2.13 | 9,590 | 831 | 1.55 | 1,400 | 1,072 |
| Pfam-Uniprot | 16,712 | 84,689,547 | 81.2 | 8.49 | 9,225 | 459 | 3.32 | 22,331 | 1,233 | 2.64 | 4,173 | 2,249 |
| Pfam-NCBI | 16,712 | 177,952,603 | 179.4 | 19.46 | 22,962 | 948 | 6.55 | 48,163 | 1,977 | 4.97 | 7,412 | 4,495 |

The top rows show the compression for a single Stockholm file for each collection. The bottom rows are for separate FASTA files (single file for each family) without metadata. File sizes are in GBs and times are in seconds. “c-time” means compression time and “d-time” means decompression time. Bold font means the best results.

Regarding compression times, MSAC is significantly faster than gzip and 7-zip in compression but slower in decompression. In real numbers, we were able to compress the 42 GB Pfam database in less than 45 minutes and decompress it in less than 15 minutes, which should be acceptable for typical scenarios.

The evaluated compressors differ significantly in their architecture, so it would be interesting to take a closer look at how they perform for protein families of different size. Therefore, we extracted all protein families and stored them in separate FASTA files. The summary of sizes and (de)compression times is given in Table 1. More details can be found in Figure 3. We grouped the FASTA files according to their sizes into disjoint subsets. The smallest size for each subset is presented at the horizontal axis, e.g., the value of 100K means that the bars above refer to files of sizes between 100KB (including) and 250KB. The height of a single bar represents the average compression ratio obtained for a given subset, while the number above the bar reflects cardinality of the subset. The largest file in this experiment was slightly smaller than 1GB. As one can see, for all size ranges (except for the smallest one containing files below 1 KB) the best compression ratio was obtained by MSAC. More importantly, the ratio rapidly increases with the size of the file. A similar trend is observed for gzip and 7zip, but their compression-ratio growth is slower.

**Fig. 3.**
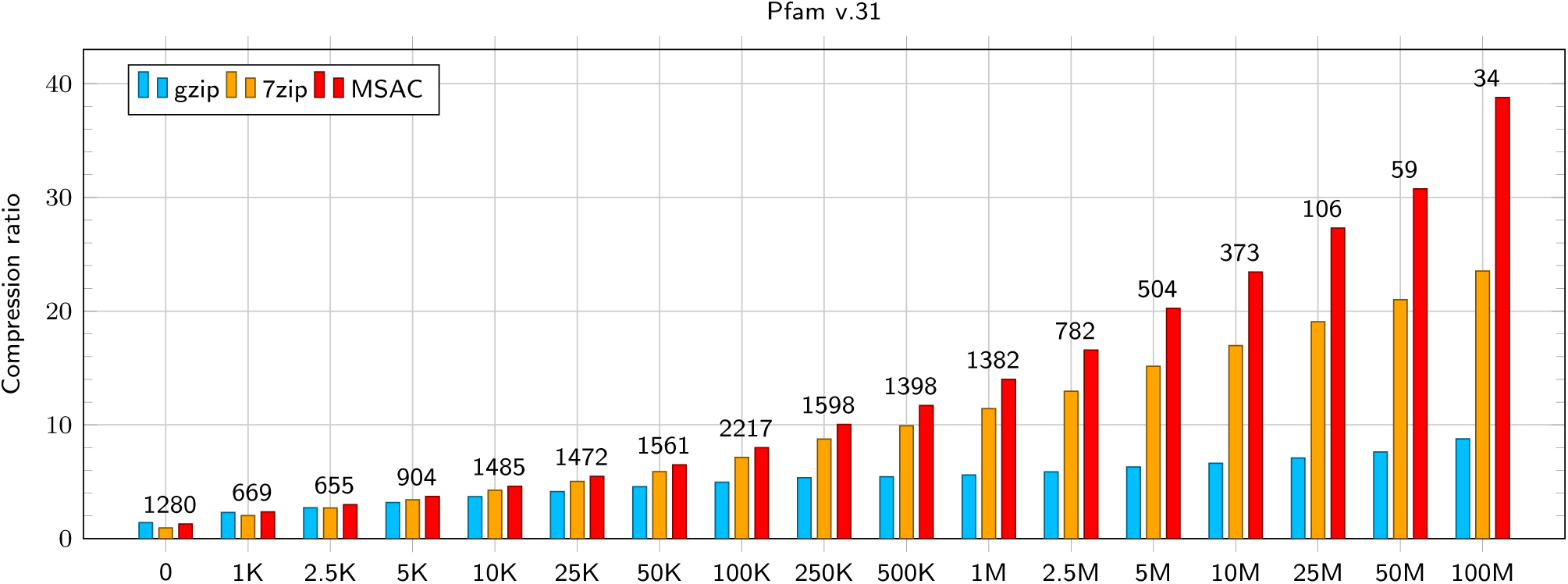
Compression ratios for subsets of Pfam collection. The sets are of given size (in bytes), e.g., the 100K set contains all files with sizes in range [100KB; 250 KB]. The number above the bars shows the number of families in a subset.

**Fig. 4.**
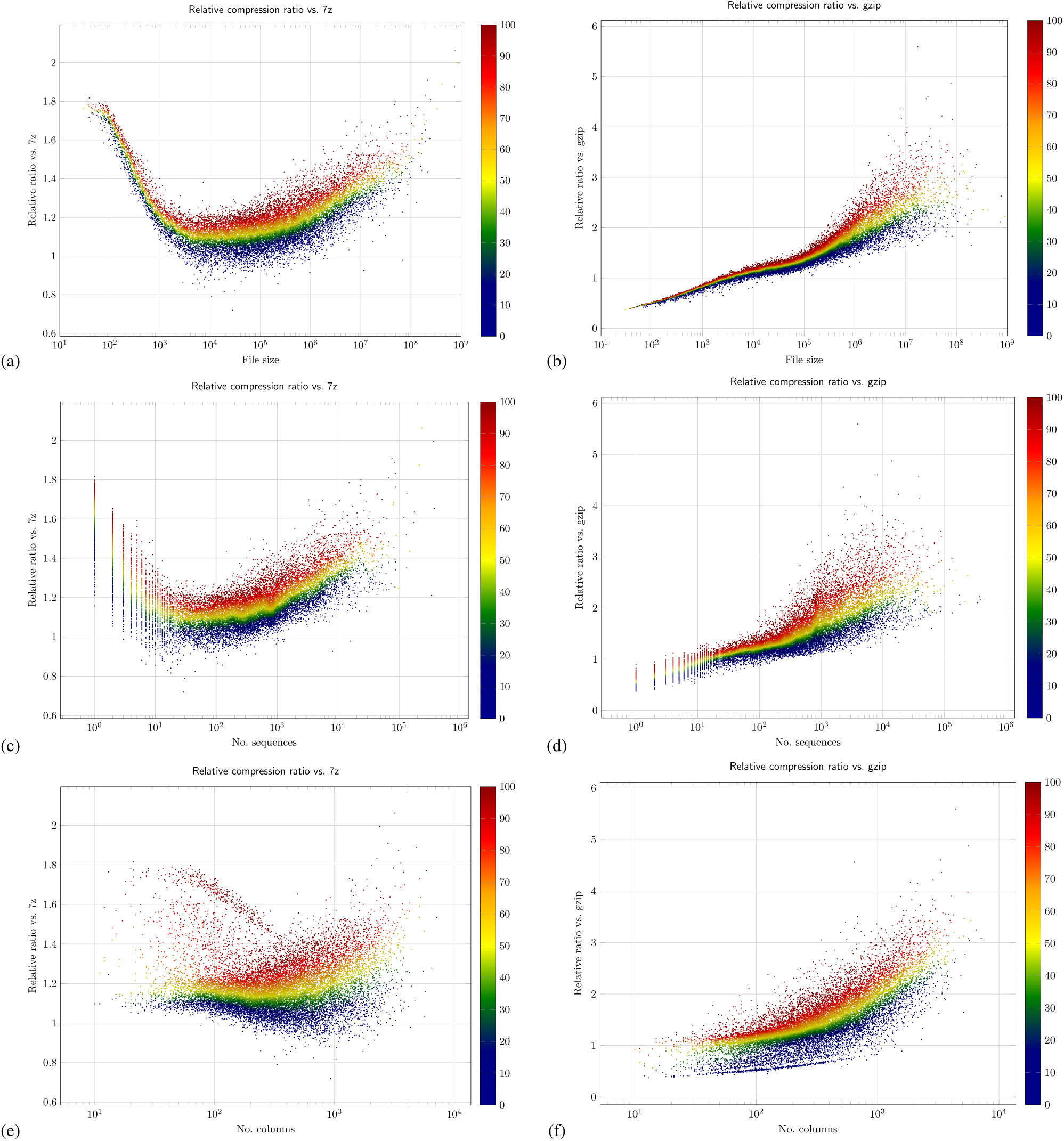
Comparison of the advantage of MSAC over 7zip (left column) and gzip (right column) compressors. The analyzes are for various: file sizes (subfigures a and b), number of sequences (subfigures c and d), and number of columns (subfigures e and f). Each point represents one of 16,479 files. The point color represent the percentile in the neighbourhood (defined as 10 percent difference in the size, sequence number, and column number, respectively).

The FASTA file size is only one of possible indicators related to MSA. We investigated also the number of sequences and the number of columns. Figure **??** shows the compression ratios for all families, i.e., each subplot contains 16,479 points. The left subplots show the advantage of MSAC over 7zip (defined as the size of 7-zip-compressed file divided by the size of MSAC-compressed file), while the right subplots show the advantage of MSAC over gzip. Since the number of points in the plots are large, for clarity we assigned a color to each of them. The color represents the rank of the advantage (over 7-zip or gzip) in group of files of comparable size (or comparable number of sequences for subfigures c and d; or comparable number of columns—subfigures e and f). By comparable we mean 10 percent neighbourhood. For example, to assign a color to some file of size 1 MB (subfigure a) we picked the advantages over 7-zip for all files up to 10 percent smaller and 10 percent larger. Then we ranked the results and calculated the percentile. Thus, the yellow color represents the median values in the neighbourhood.

As one can observe, all medians are always above 1.0 for 7-zip and there is a growing tendency in the advantage for very small and very large files. Similar observations can be made, when the size of the family is defined as the number of sequences (subfigure c), but for growing number of columns (subfigure e) the trend is less clear. The maximal advantage over gzip is much larger in real numbers (up to 7.5, compared to 2.1 for 7-zip). For tiny files (smaller than 300 bytes), the median of the advantage is below 1.0, which means that gzip performs better. Nevertheless, compressing of such small files is considered to be useless, as the gains are negligible.

It is also noteworthy to say that the largest single protein family from Pfam-NCBI (PF07690.15) consumes about 4.36 GB of space. MSAC was able to reduce this to 81.7MB (150.2MB for 7-zip nad 543.5MB for pigz). The MSAC compression and decompression times were 120 s and 95 s, respectively (791 s and 33 s for 7-zip; 778 s and 13 s for pigz). As the modern algorithms for MSA determination, like MAFFT (Katoh & Standley, 2013), Clustal Omega (Sievers *et al*., 2011), PASTA (Mirarab *et al*., 2015), FAMSA (Deorowicz *et al*., 2016), to name a few, are able to process families containing more than 100,000 sequences in a few hours (or even less) at modern workstations, and the resulting alignments sometimes consume tens of GB, the ability of MSAC to significantly reduce the necessary space in a very-short time is remarkable.

## 4 Discussion

The Pfam database is a growing collection of protein families. The raw data for its default variant occupy about 42 GB, which causes problems with both storage and data transfer. Currently, the most popular method to reduce the file sizes is the commonly known, universal compressor—gzip program. When applied to the basic Pfam dataset it allows to reduce the space to about 5.6GB. From the practical point of view, application of gzip still leads to important gains, but we can be almost sure that the size of the discussed collection will be growing in the near future. Even now, the largest variant of Pfam v. 31.0 consumes ~230GB for raw data and ~30GB, when gzip-compressed.

In this article, we proposed a novel compression algorithm for multiple sequence alignments files. The algorithm works in a few stages. A generalisation of the positional Burrows–Wheeler transform for non-binary alphabets is performed as the first stage. Then, transforms adopted from the literature are determined, namely weighted frequency counter transform (WFC) followed by ran length encoding transform (RLE-0). Finally, the modelling stage for the entropy coder is executed.

The experiments made for three variants of the largest existing protein family database, Pfam, showed that MSAC offers on average more than 3 times better compression ratio than gzip for the whole collection of families. The advantage over 7-zip is smaller, but still significant. What is important, MSAC advantage over gzip and 7zip grows for growing family size, which is promising, as in the future the protein families will be definitely larger. Even today, it seems that the very good compression ratio and good compression and decompression speeds allows to consider MSAC as a replacement for gzip for those that suffers from huge size of MSA collections.

## Acknowledgements

The work was supported by Polish National Science Centre under the project DEC-2015/17/B/ST6/01890 and performed using the infrastructure supported by POIG.02.03.01–24–099/13 grant: ‘GeCONiI—Upper Silesian Center for Computational Science and Engineering’.

